# Complex organic matter degradation by secondary consumers in chemolithoautotrophy-based subsurface geothermal ecosystems

**DOI:** 10.1101/2023.01.20.524876

**Authors:** Raegan Paul, Timothy J. Rogers, Kate M. Fullerton, Matteo Selci, Martina Cascone, Murray H. Stokes, Andrew D. Steen, J. Maarten de Moor, Agostina Chiodi, Andri Stefánsson, Sæmundur A. Halldórsson, Carlos J. Ramirez, Gerdhard L. Jessen, Peter H. Barry, Angelina Cordone, Donato Giovannelli, Karen G. Lloyd

## Abstract

Microbial communities in terrestrial geothermal systems often contain chemolithoautotrophs with well-characterized distributions and metabolic capabilities. However, the extent to which organic matter produced by these chemolithoautotrophs supports heterotrophs remains largely unknown. Here we compared the abundance and activity of peptidases and carbohydrate active enzymes (CAZymes) that are predicted to be extracellular identified in metagenomic assemblies from 63 springs in the Central American and the Andean convergent margin (Argentinian backarc of the Central Volcanic Zone), as well as the plume-influenced spreading center in Iceland. All assemblies contain two orders of magnitude more peptidases than CAZymes, suggesting that the microorganisms more often use proteins for their carbon and/or nitrogen acquisition instead of complex sugars. The CAZy families in highest abundance are GH23 and CBM50, and the most abundant peptidase families are M23 and C26, all four of which degrade peptidoglycan found in bacterial cells. This implies that the heterotrophic community relies on autochthonous dead cell biomass, rather than allochthonous plant matter, for organic material. Enzymes involved in the degradation of cyanobacterial- and algal-derived compounds are in lower abundance at every site, with volcanic sites having more enzymes degrading cyanobacterial compounds and non-volcanic sites having more enzymes degrading algal compounds. Activity assays showed that many of these enzyme classes are active in these samples. High temperature sites (> 80°C) had similar extracellular carbon-degrading enzymes regardless of their province, suggesting a less well-developed population of secondary consumers at these sites, possibly connected with the limited extent of the subsurface biosphere in these high temperature sites. We conclude that in < 80°C springs, chemolithoautotrophic production supports heterotrophs capable of degrading a wide range of organic compounds that do not vary by geological province, even though the taxonomic and respiratory repertoire of chemolithoautotrophs and heterotrophs differ greatly across these regions.

## Introduction

Terrestrial geothermal systems emit volatiles from the Earth’s interior (i.e., mantle and crust) to the atmosphere. Often, meteoric water permeates into the subsurface hydrothermal system, where it is heated and rises to the surface, bringing with it volatiles from the deep subsurface [1]. The differential enrichment of these volatiles into geothermal fluids creates environmental niches that can be saturated with deeply-derived inorganic carbon and other compounds [2–4]. In convergent margins such as those of the Central American and the Andean Central Volcanic Zone, inorganic carbon is derived from the mantle, overlying crust and/or down-going slab [5]. In divergent spreading centers and/or areas with mantle plume-influenced volcanism, such as Iceland, geothermal systems are often dominated by deep mantle gases (e.g., Hardardottir et al. 2018 [6]). These geological systems create a large diversity of surface-emitting springs that range in temperature, pH, inorganic carbon content, and availability of redox active compounds that together make the driving force of microbial community composition [4,7,8]. Sampling fluid emissions from natural surface springs provides access to these deeply-sourced microbial communities and the volatiles that support them [8–11].

The important role that chemolithoautotrophs play in these geothermal ecosystems is well-established (e.g., [4,12–14]). The heterotrophic communities within these systems are less often studied, even though heterotrophs have been shown to be dominant within heavily-sedimented subsurface ecosystems (e.g., [15]). These heavily-sedimented systems do not have a constant supply of redox active volatiles and are therefore dependent on allocthonously-derived organic matter, and likely differ greatly from geothermal ecosystems. Recent work has focused on understanding the heterotrophic community within terrestrial geothermal systems [16,17] and many industrially useful carbohydrate- and peptide-degrading enzymes have been isolated from these microbial communities [18,19]. The taxonomy and respiratory pathways of primary producers and heterotrophs are known to vary along geological gradients according to changes in deep volatile delivery [4,7,8]. However, it is not known whether organic carbon degradation pathways vary along with them. Different types of chemolithoautotrophs may promote different compositions of carbon-degrading enzymes in the ecosystems they support. Thus, it is important to investigate the heterotrophic community because it actively participates in the geochemical cycle of terrestrial geothermal environments by consuming organic carbon and releasing inorganic carbon.

Hot springs typically contain little photosynthetically-derived organic matter, potentially leading heterotrophs to depend primarily on the byproducts of the chemolithoautotrophic community [20]. To access these organic byproducts, heterotrophs use extracellular enzymes to break down larger organic molecules into smaller molecules that can permeate their cell membrane more readily [21] ultimately recycling carbon through this autotroph-heterotroph mutualism. The composition of carbon-degrading enzymes may therefore show whether chemolithoautotrophy or photosynthesis is more important for heterotrophic communities. Here, we focus on two broad classes of extracellular enzymes: peptidases which break down proteins and carbohydrate-active enzymes (CAZymes) which break down polysaccharides and related macromolecules.

We compare springs across the Central American and the Andean (Argentinian backarc of the Central Volcanic Zone) convergent margins as well as the plume-influenced Iceland plate boundary. These regions are defined by their position across the convergent margin or continental intra-plate setting toward an oceanic plate boundary. The Costa Rican subduction zone is driven by the Cocos-Nazca plate subducting under the Caribbean plate. Northern Costa Rica is characterized by having higher volcanic activity than the other areas [22]. The Panama slab window is a result of a tear within the Nazca plate where arc volcanism ceases [23]. The formation of the slab window is also responsible for the cease in volcanism within the Cordillera Talamanca region [24], and a change in the rock chemistry in the area that shows hot spot-like compositions ([25]). The Andean convergent margin is driven by the Nazca plate subducting under the South American plate ([26]), whereas Iceland is associated with spreading along the Mid-Atlantic Ridge under the influence of a mantle plume (e.g., [2,27,28]). Environmental factors also vary significantly between the sites due to large variations in latitude, elevation, and rainfall. This wide range of geological and environmental settings provides an opportunity to study geochemically diverse springs. Sites from Argentina and Iceland were placed into their own categories because they have different tectonic processes than those of Central America. The sites are then color coded based on these different geological processes that distinguish them (Figure 1).

**Figure 1.**
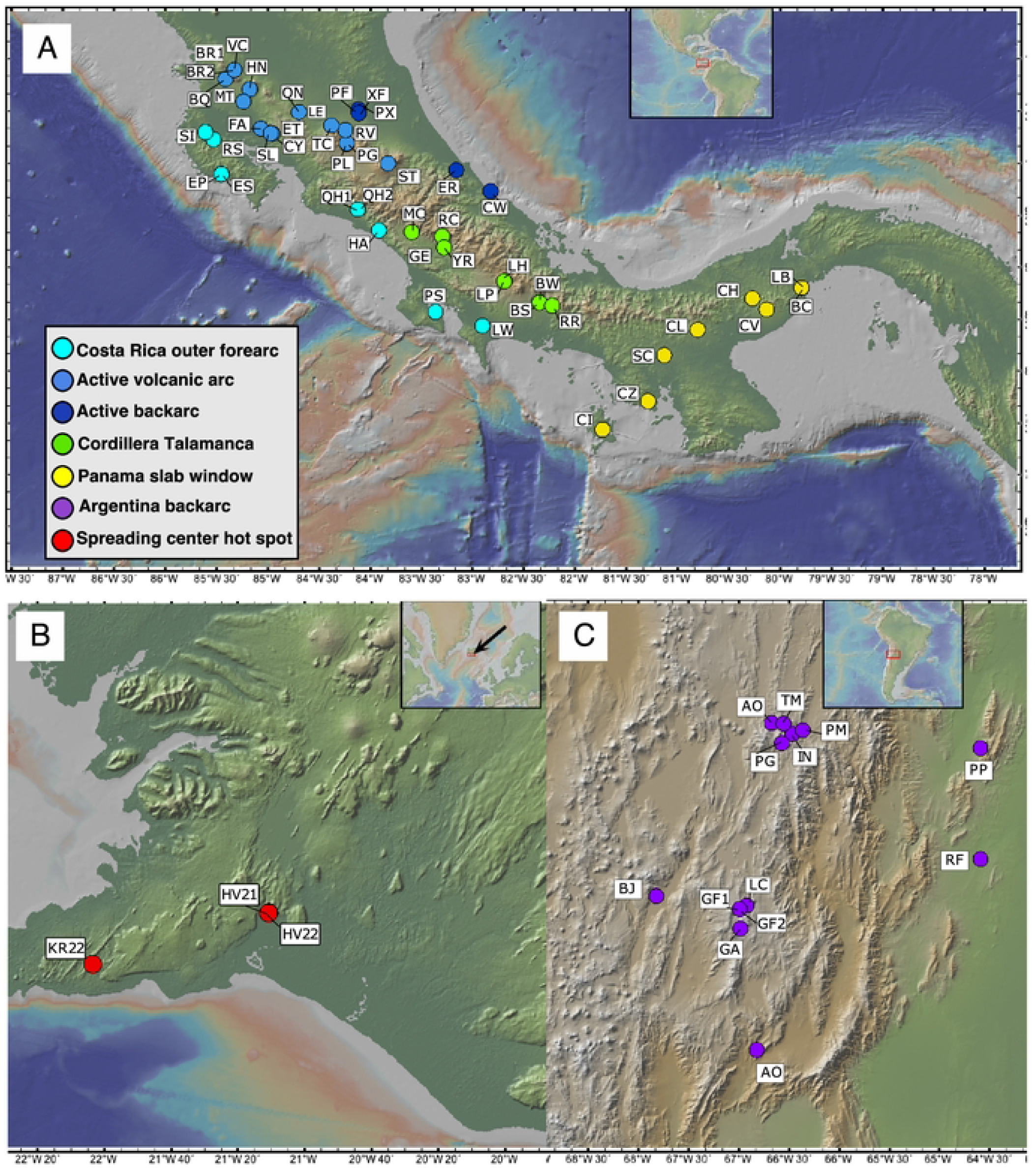
Maps of site locations. **A**. Costa Rica and Panama sites are color coded to match the different geological provinces. **B.** Argentina sites are all backarc. C. Iceland sites are all active volcanic hot spot and spreading center.

Enzyme assays and bioinformatics analyses on metagenomes were used to analyze the microbial community interactions within diverse geothermal systems. To study the heterotrophic activity within geothermal systems, assemblies from the Costa Rican convergent margin, the Argentina backarc of the Andean convergent margin and the subaerial section of the Mid-Atlantic ridge (Iceland) were annotated using dbcan2 to find CAZymes and DRAM to annotate the MEROPS families [29,30]. MEROPS classifies proteolytic enzymes using hierarchical classification by homologous sequences [31]. The CAZyme database splits carbohydrate-active enzymes into classes of glycoside hydrolases (GH), glycosyl transferases (GT), polysaccharide lyases (PL), carbohydrate esterases (CE), auxiliary activities (AA), and carbohydrate binding molecules (CBM), defined by sequence similarity [32]. Enzyme commission numbers are based on the reactions catalyzed instead of sequence homology [33]. Using these different tools allows for a broad analysis of the potential organic matter degrading functions of proteins based on their sequence homology. Hierarchical clustering and principal coordinate analysis were used to find correlations between the sites and the enzyme families found within them. By combining the maximum potential enzymatic activity, measured by low molecular mass fluorogenic substrate proxies, with the metagenomic annotations, we can see a larger scope of the potential heterotrophic activity within these sites.

## Methods

### Sampling

Sample collection was performed following the protocols previously described [3,4,7] and using the rationale described by Giovannelli et al. 2022 [11]. Sites are color coded based on province (Figure 1). GPS coordinates, site names, temperature and pH measurements are shown in Table 1. From each site, temperature and pH were measured directly in the fluids using a portable YSI Plus 6-Series Sonde Multimeter (YSI Incorporated, Yellow springs, OH) and 0.5 to 1.5 liters of hydrothermal fluids venting from the subsurface were collected. Care was taken when collecting fluids to do this as close to the perceived fluid source as possible. Fluids were immediately filtered through Sterivex 0.22 μm filter cartridges (MilliporeSigma) and quick-frozen onsite in a liquid nitrogen-cooled dry shipper. When fluid sampling was complete and to avoid resuspension, ~10 mL of surface sediments constantly overwashed by the venting source were placed into a sterile plastic vial and frozen onsite along with the filters. Sample names ending in (f) are from filtered fluids and those ending in (s) are from surface sediments.

**Table 1:**
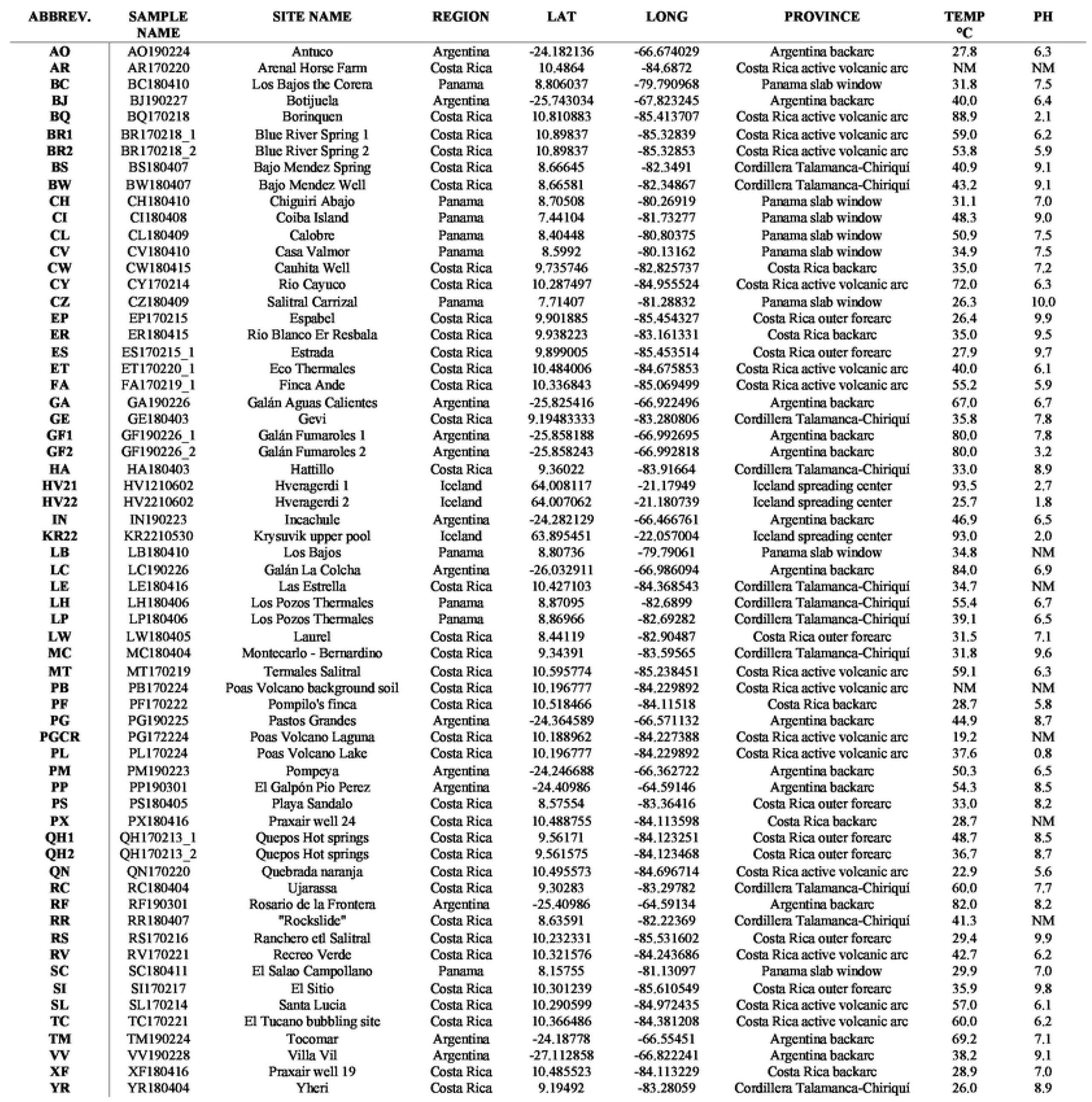
Site name and abbreviation supplemented with province, latitude, longitude, temperature, and pH. NM means not measured.

### Bioinformatic Processing

For the Iceland metagenomes, raw reads were trimmed with Trimmomatic (v 0.39) [34] and assembled using the MetaWRAP (v 1.3.2) pipeline [35]. The quality of the Iceland assemblies was determined using Quast (v4.4) [36] on Kbase [37]). All other assemblies were generated by trimming raw reads and mate-pairing them with Trimmomatic (v 0.38). Reads were assembled *de novo* with metaSPAdes with a minimum contig length of 1.5 kb [7]. Assemblies LCf, LCs, RFs, and TMs were assembled using MEGAHIT (v 1.2.9) [38]. The assemblies were annotated using prokka (v 1.14.5) [39] and dbcan2 [29]. For peptidase annotations the assemblies were uploaded to KBase [37] and annotated with DRAM (v.0.1.0) [30]. Secreted proteins were identified using SignalP (v 2.0) [40]. The clean reads were mapped back to the assemblies for read coverage using bowtie2 (v 2.3.5.1) [41]. Hierarchical clustering based on spearman correlation, was performed using hclust from the base R package (R version 4.2.1). Total microbial community analysis of these sites is the focus of previous work, therefore, taxonomic identification was only performed for contigs for the high temperature sites HV21, HV22, and KR22 using gottcha2 on Kbase (Figure S1) [42].

CAZy family abundances were annotated using dbcan2. Annotations were selected if they were annotated with at least two tools: HMMER and DIAMOND. The annotated gene IDs were then combined with the prokka gene ID and contigs to gain the abundance of each contig. Then the read abundance of each annotation normalized to the total assembly size was used to generate a heatmap with hierarchical clustering of the CAZyme families and site locations. All annotations that are presented also were annotated for having a signal peptide sequence by SignalP [40]. Protein annotations that are not present within 75% of the assemblies were removed for better visualization.

Enzyme commission numbers were annotated with dbcan2 and combined with the SignalP annotations to estimate secretion. Enzyme commission groupings are only shown for class 3 hydrolases where the enzyme commission number was present in at least 75% of the assemblies. MEROPS peptidase annotations were completed using DRAM on Kbase and annotated using SignalP. The gene IDs were then matched to contigs of the assemblies to get the normalized read coverage.

### Enzymatic Assays

Enzymatic assays were performed following the methods of Bell (2013) [43] with slight modifications. Briefly, sediments were weighed out at 2.75 grams wet weight. The sediments were then combined with 91 mL of 0.5 M Tris-HCl buffer with a matching pH to the original site locations. The sediment slurries were blended for one minute to homogenize them. Then 800 μL of each slurry was pipetted into deep well plates in duplicate. 200 μL of substrates with a concentration of 200 μM were then added to each well. The substrates used were 4-Methylumbelliferyl α-D-glucopyranoside (AG), 4-Methylumbelliferyl β-D-glucopyranoside (BG), 4-Methylumbelliferyl β-D-cellobiosidase (CB), 4-Methylumbelliferyl N-acetyl-β-d-glucosaminidase (NAG), L-Leucine-7-amido-4-methylcoumarin (LEU), 4-Methylumbelliferyl phosphate (PHOS), 4-Methylumbelliferyl sulfate potassium salt (SULF), 4-Methylumbelliferyl-β-D-xylopyranoside (XYL). Each deep well plate was incubated for 3 hours at 30-70 °C, and time points were taken at 0 hours, 1.5 hours, and 3 hours. To take each time point, 200 μL was pipetted from the deep well plate to a black flat bottom 96 well plate. The fluorescence was measured at 455 nm after excitation at 355 nm using a Tecan Infinite M200 Pro Fluorimeter. Differences in enzyme activities among provinces were tested using a Kruskal-Wallis test, implemented in R, due to the strong non-normal distribution of activities in the data set.

## Results

### Sites

Three different geographical areas were analyzed: the Central American and Andean convergent margins, and Iceland (mantle plume-influenced spreading center). Large variations in fluid sources (i.e. mantle, slab, crust, or surficial) are expected due to the contrasting geologic and environmental settings of the studied areas. We include data from 22 sites in Costa Rica that are influenced by the convergent margin, with analyses of the geochemistry, respirations and taxonomic identities published previously [3,4,7,44]. We include 47 additional sites spanning Costa Rica and Panama that are also influenced by the convergent margin, 13 sites from the Andean convergent margin and 3 sites from Iceland. Sites were grouped by geographic-tectonic setting: Costa Rica outer forearc, Costa Rica active volcanic arc, Costa Rica backarc, Cordillera Talamanca, Panama, Argentina backarc, and Iceland (Figure 1, Table 1). These groupings allow us to explore large variations in fluid sources and physicochemical characteristics which are ultimately related to their tectonic setting. For example, in Costa Rica the distance to the trench (outer forearc to active arc to backarc) correlates with large variations in temperature, pH, and mantle-derived components that affect chemical compositions of the fluids and the microbial communities [4,45]. The sites with the lowest temperatures (22.9 – 29.9 °C) are QN, YR, CZ, EP, AO, ES, PF, RS, and SC (Table 1). The sites with the highest temperatures (80-93.5 °C) are GF1, GF2, RF, LC, BQ, KR22, HV21 and HV22. Sites with no recorded temperature were AR and PB, which are also background sediments near volcanic soils. Sites that had the lowest pH (0.85 – 3.21) were PL, HV21, HV22, KR22, BQ, and GF2. Sites with the highest pH (9.0 – 10.0) were CZ, EP, RS, SI, ES, MC, VV, BW, BS, and CI. Sites without a recorded pH were LB, RR, AR, LE, PB, PGcr, and PX.

### Enzyme Activities

Most of the hydrolysis rates measured were indistinguishable from zero (Figure 2). This could mean either that few of the enzymes were being expressed at the time of sampling or that we were unable to properly recreate the geochemical conditions present *in situ*. Of the enzymes tested, carbon-acquiring enzymes (AG, BG, CB, XYL) were more active than enzymes associated with phosphorus and sulfur acquisition (PHOS and SULF) (Table S3). LEU hydrolysis was orders of magnitude lower than the other enzymes assayed, in contrast to soils where LEU often has high hydrolysis rates [46]. NAG hydrolysis was positive at more sites than the other substrates. PHOS hydrolysis had only one site with zero activity, CL. From the Kruskal-Wallis statistical analysis, no significant differences (p>0.05) were observed for any enzyme activities between provinces.

**Figure 2:**
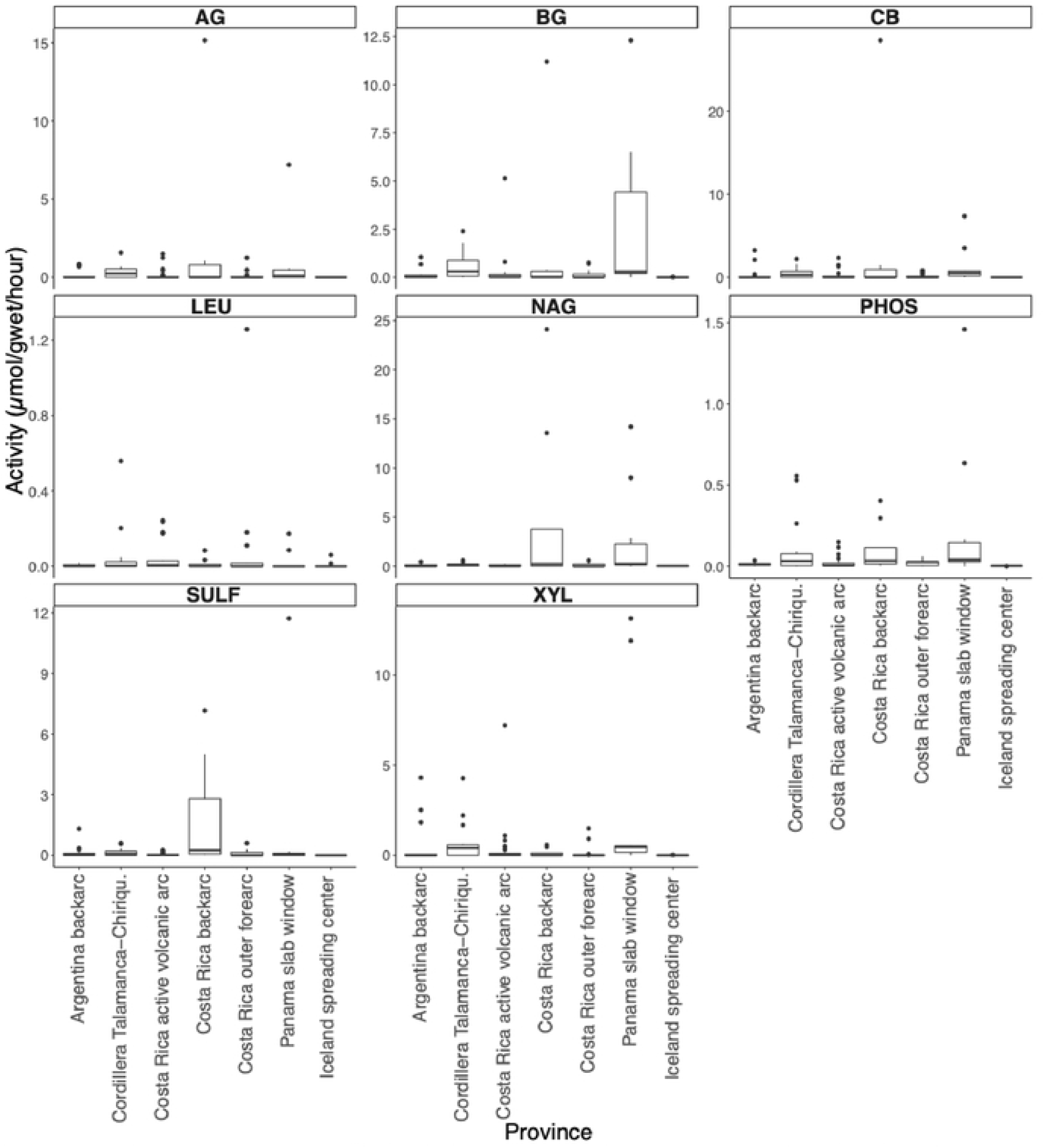
Enzyme activity box plot. Activities for each site were grouped together based on province. Activity measurements were only performed on sediment samples. Each activity is presented in pmol/g_wet_/h. Enzyme substrates were separated into different panels. Enzymes abbreviations are alpha-glucosidase (AG), beta-glucosidase (BG), cellobiohydrolase (CB), leucine aminopeptidase (LEU), N-acetyl-β-D-glucosaminidase (NAG), phosphatase (PHOS), sulfatase (SULF), and xylosidase (XYL).

### Metagenomic assemblies

Of the three metagenomic assemblies from Iceland, HV22 has 4,344 contigs, KR22 has 3,214, and HV21 has 912 (Table S1). The total length of these assemblies is 5,645,364 bp for HV21, 7,802,221 bp for HV22, and 12,561,178 bp for KR22. The total contig numbers for the Costa Rica, Panama and Argentina assemblies range from 488 to 164,798 (Table S2). The total sizes of the Costa Rica, Panama and Argentina assemblies range from 8,678,655 to 509,373,734 (Table S2). At the genus level, HV21 has 70% of reads annotated as *Sulfolobus*, 15% as *Acidianus*, and the 8% as *Thermoproteus*. The remaining reads are distributed across *Methylorubrum, Pseudomonas*, and *Stenotrophomonas*. HV22 had 38% of reads annotated as *Sulfolobus*, 21% as *Thermoproteus*, 10% as *Cutibacterium*, and the rest are distributed across other genera. KR22 has 45% of reads identified as *Acidianus*, 8% as *Sulfolobus*, and 5% as *Thermoproteus*, with the remaining taxonomy distributed across other genera.

### Peptidases

In total, there are 144 MEROPS family annotations predicted to be secreted (Figure 3), M (Metallo) with 59, S (Serine) with 32, C (Cysteine) with 29, A (Aspartic) with 9, N (Asparagine) with 5, T (Threonine) with 4, U (Unknown) with 4, G (Glutamic) with 1, and P (Mixed) with 1 (Figure 3). Of these 144 families, 88 are present in every assembly. The total read abundance of annotations for MEROPS is 283,488,080, which is much greater than those of CAZy at 1,729,668, and EC3 at 633,511, which are discussed below.

**Figure 3:**
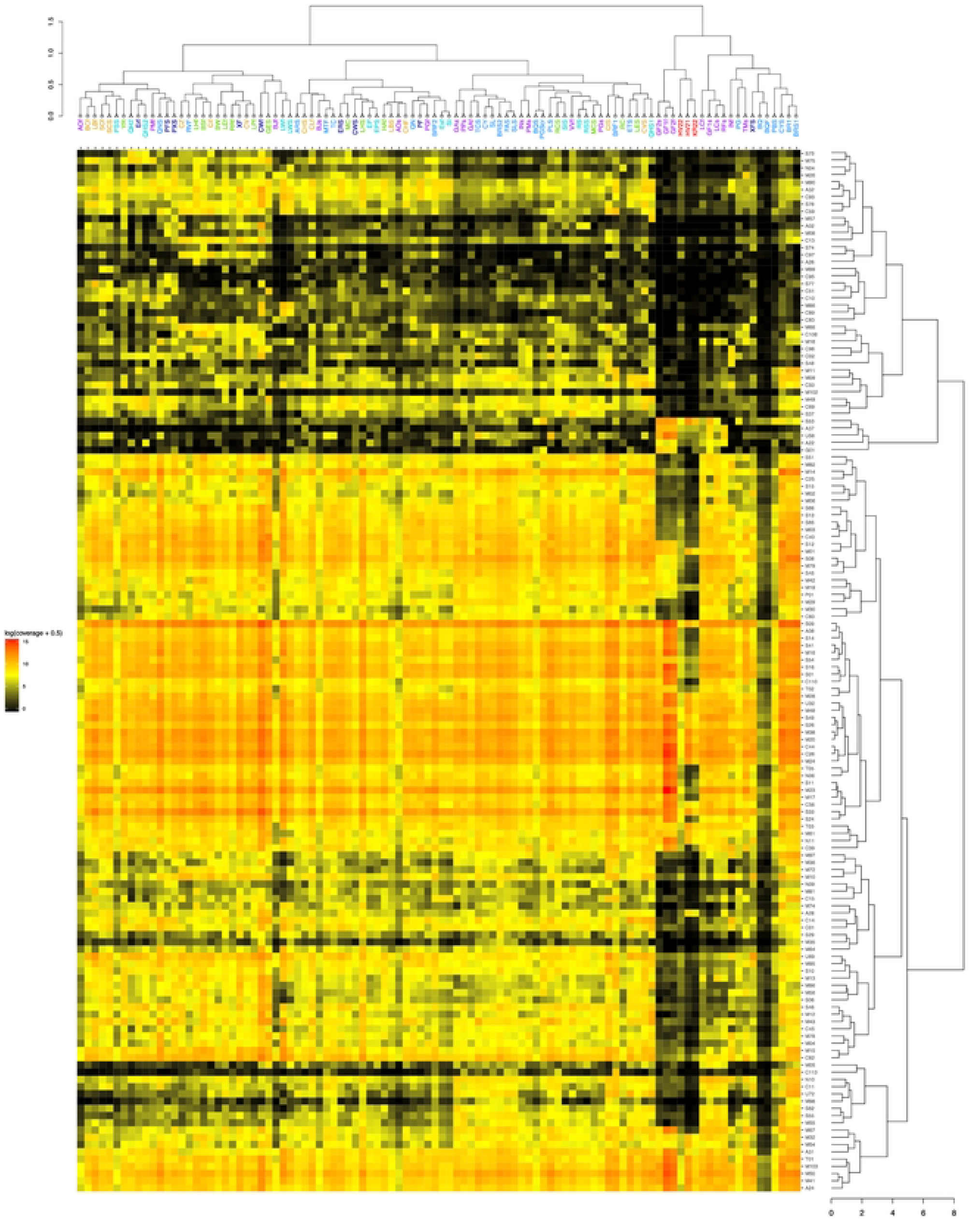
Heatmap with hierarchical correlation of read abundances of MEROPS families (y axis) per site (x axis) based on spearman rank correlation. Abundance shown in log of normalized contig abundance with 0.5 added to avoid zeros for visualization. Untransformed abundances were used for the spearman correlation. Sites are colored by the geological provinces shown in Figure 1.

The highest read abundance normalized to total assembly size is 5,316,108.99 for M23 in site GF1f. The most abundant MEROPS normalized read abundances are S09 (prolyl oligopeptidase), M23 (beta-lytic metallopeptidase), C26 (gamma-glutamyl hydrolase), S33 (prolyl aminopeptidase), M38 (isoaspartyl dipeptidase), C44 (amidophosphoribosyl transferase precursor), S49 (signal peptide peptidase A), M20 (glutamate carboxypeptidase), M50 (site 2 peptidase), and S16 (Lon-A peptidase). The sites with the least amount of MEROPS annotations are GF2f, HV21, GF2s, and GF1f.

Within the spearman correlation hierarchical clustering of sites based on the MEROPS families, the highest temperature sites (GF2s, GF1f, GF2f, HV22, HV21, and KR22) cluster together. Some sites’ sediment and fluid assemblies cluster together: SC, LW, EP, RS, and BR1. One cluster consists of only fluid samples CZ, RVF, LHf, BSf, Cif, BW, LEf, RRf, XF, CV, and LPf. For the hierarchical clustering of MEROPS families based on their distribution across sites, one cluster contains MEROPS families S53, A37, U56, A22, G01, that are highly present in only the very hot sites. It has been shown that these peptidase families are associated with acidophilic or thermophilic archaea [47].

### CAZymes

CAZy families predicted to be secreted are present in all sites at orders of magnitude lower read abundance than peptidases. Of the six CAZy classes, the most abundant are the glycosyl hydrolases (GH). CAZy families that are abundant in all sites are GH23 (peptidoglycan lyase), CBM50 (binds peptidoglycan and chitin), GH102 (peptidoglycan lytic transglycosylase), and GH103 (peptidoglycan lytic transglycosylase). All five CAZy classes are present in 76 of the assemblies with 89 annotations from glycoside hydrolases (GH), 31 from carbohydrate binding modules (CBM), 17 from polysaccharide lyases (PL), 10 from carbohydrate esterases (CE), 7 from glycosyltransferases (GT), and 2 from auxiliary activities (AA). There are a total of 156 CAZy families present. The assemblies with the least number of CAZyme annotations are HV21, KR22, GF1f, GF2f, GF2s and BQf. Sites that contain less than 75% of the annotated CAZyme families are BQ, RFs, BQF, LCs, GF2s, HV21, KR22, GF2f, and GF1f; all of which had temperatures over 80°C.

Within the spearman correlation of the site clustering, we see a grouping of sites that are from the Argentina backarc and have a higher temperature range (Figure 4). These sites cluster together due to the low abundance of enzyme annotations within their assemblies. Some sites’ sediment and fluid samples cluster together such as, LB, SC, BJ, QN, AO, RS, QH2, ER, and LC.

**Figure 4:**
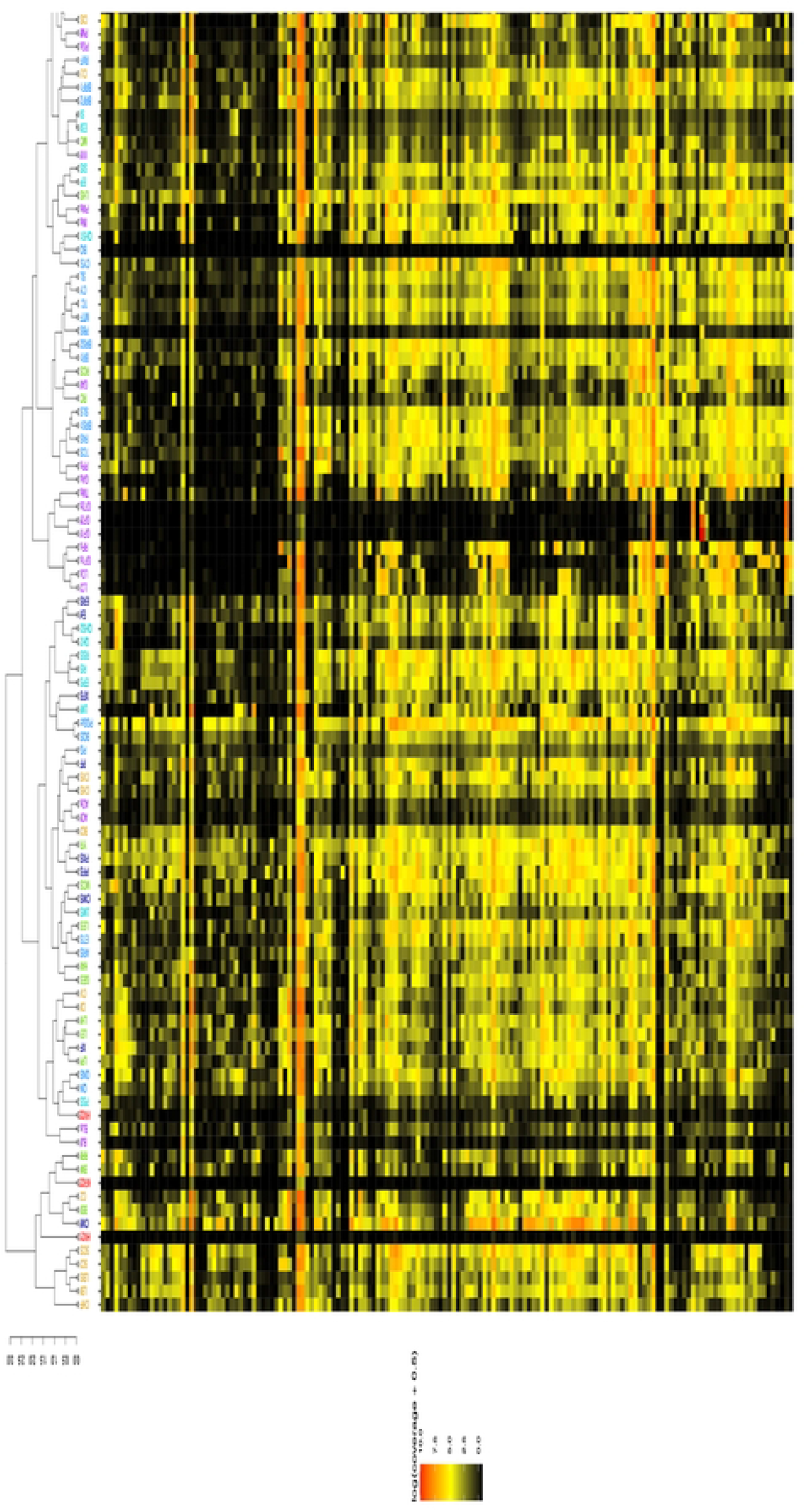
Heatmap with hierarchical correlation of read abundance of CAZy families (y axis) per site (x axis) based on spearman rank correlation. Visualization details are the same as Figure 3.

### Enzyme Commission

Enzymes from EC3 are hydrolases, so many of them overlap with those in the CAZyme and MEROPS groups, but the EC categories are more finely divided than CAZyme groups, so they describe hydrolase functionality more precisely. As with CAZymes, EC3 annotations have orders of magnitude lower read abundance than the MEROPs peptidases (Figure 5). There are a total of 117 EC hydrolase annotations. The EC hydrolases are subdivided into 3.1 (ester hydrolases), 3.2 (glycosylases), 3.3 (ether hydrolases), 3.4 (peptidases), 3.5 (other non-peptide carbon and nitrogen hydrolases), 3.6 (acid anhydride hydrolases), 3.7 (other carbon-carbon bond hydrolases), 3.8 (halide hydrolases), 3.9 (other phosphorus nitrogen bond hydrolases), 3.10 (sulfur nitrogen bond hydrolases), 3.11 (other carbon phosphorus bond hydrolases), 3.12 (sulfur-sulfur bond hydrolases), 3.13 (carbon sulfur bond hydrolases). The spearman site correlation shows a clustering of sites with very few hydrolases present (Figure 5). The sites that have less than 75% of the protein annotations shown are INf, LCf, HV22, RFs, BQ, GF1s, BQF, KR22, HV21, LCs, GF2s, GF2f, and GF1f. All these sites except for IN fall within the highest temperature range (80-93.5°C). The most abundant EC annotation is 3.4.21.107 (peptidase Do). Other ECs in high abundance are N-acetylmuramoyl-L-alanine amidase (3.5.1.28), subtilisin (3.4.21.62), C-terminal processing peptidase (3.4.21.102), prolyl oligopeptidase (3.4.21.26), oryzin (3.4.21.63), triacylglycerol lipase (3.1.1.3), and beta-lactamase (3.5.2.6). The most common category among these high abundance enzymes is EC group 3.4, which are peptidases.

**Figure 5:**
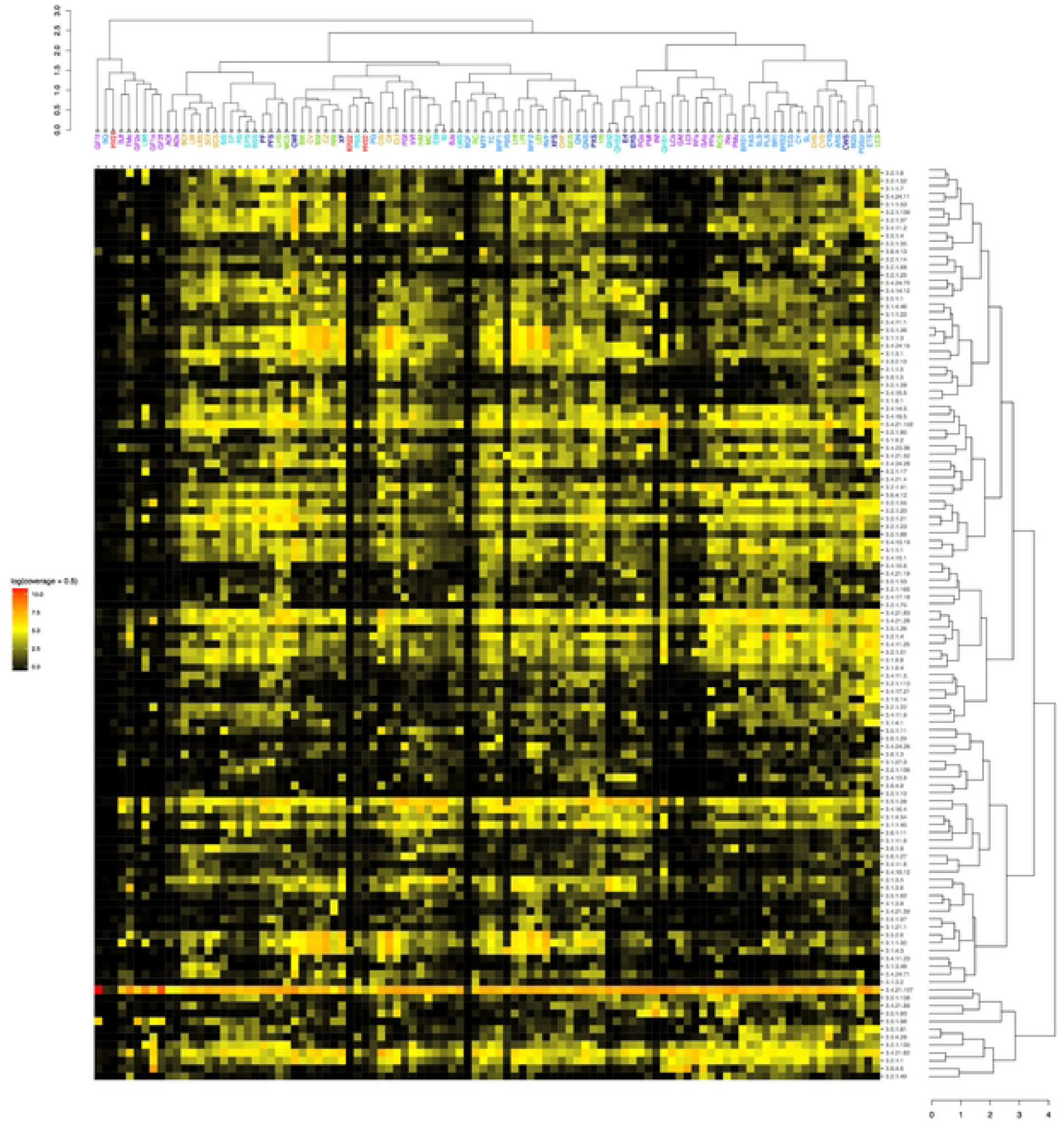
Heatmap with hierarchical correlation of read abundance of hydrolases (EC 3, y axis) per site, based on spearman rank correlation. Visualization details are the same as Figure 3.

The EC numbers that are present in all assemblies are alkaline phosphatase (3.1.3.1), (3.2.1.1), beta-glucosidase (3.2.1.21), alpha-L-fucosidase (3.2.1.51), dipeptidyl-peptidase IV (3.4.14.5), peptidyl-dipeptidase A (3.4.15.1), peptidyl-dipeptidase Dcp (3.4.15.5), peptidase Do (3.4.21.107), subtilisin (3.4.21.62), oryzin (3.4.21.63), and endothelin-converting enzyme 1 (3.4.24.71). Three of the EC numbers that are present in all assemblies are also some of the most abundant (3.4.21.107, 3.4.21.62, and 3.4.21.63). Of the eleven EC numbers present in all assemblies, seven are within the family of peptidases. As with the MEROPS and CAZymes annotations, some of the fluid and sediments from the same site group together: AO, LB, SC, PF, CI, QN, QH2, and ER.

### CAZy degradation groupings

We summed the abundance of CAZy families that are either part of chitin/peptidoglycan degradation (bacterial necromass) or xylan degradation (photosynthate products) (Tables 3 and 4). In total cell-degrading enzymes are in higher read abundance (521,794.37) than photosynthate-degrading enzymes (227,434.98) (Table 4). The assemblies with the highest percentage of cell-degrading enzymes are BJf, SI, KR22, ESf, RVf, and CIf. Only 20 assemblies have fewer than 50% of the annotations for cell-degrading enzymes. The abundance of these enzymes for each site, with sediment and fluid components, is shown in a stacked bar plot to visualize the differing quantity across sediments and fluids (Figure 6). Overall, cell-degrading enzymes are more abundant than photosynthate-degrading enzymes. Photosynthate-degrading enzymes are more abundant in sediments than in fluids, whereas the cell-degrading enzymes are more abundant in fluids than in sediments (Table 4). CAZy family GH103 is more abundant in fluids than in sediments (Figure 6). CAZy families GH18 and GH3 are more abundant in sediments than fluids.

**Figure 6:**
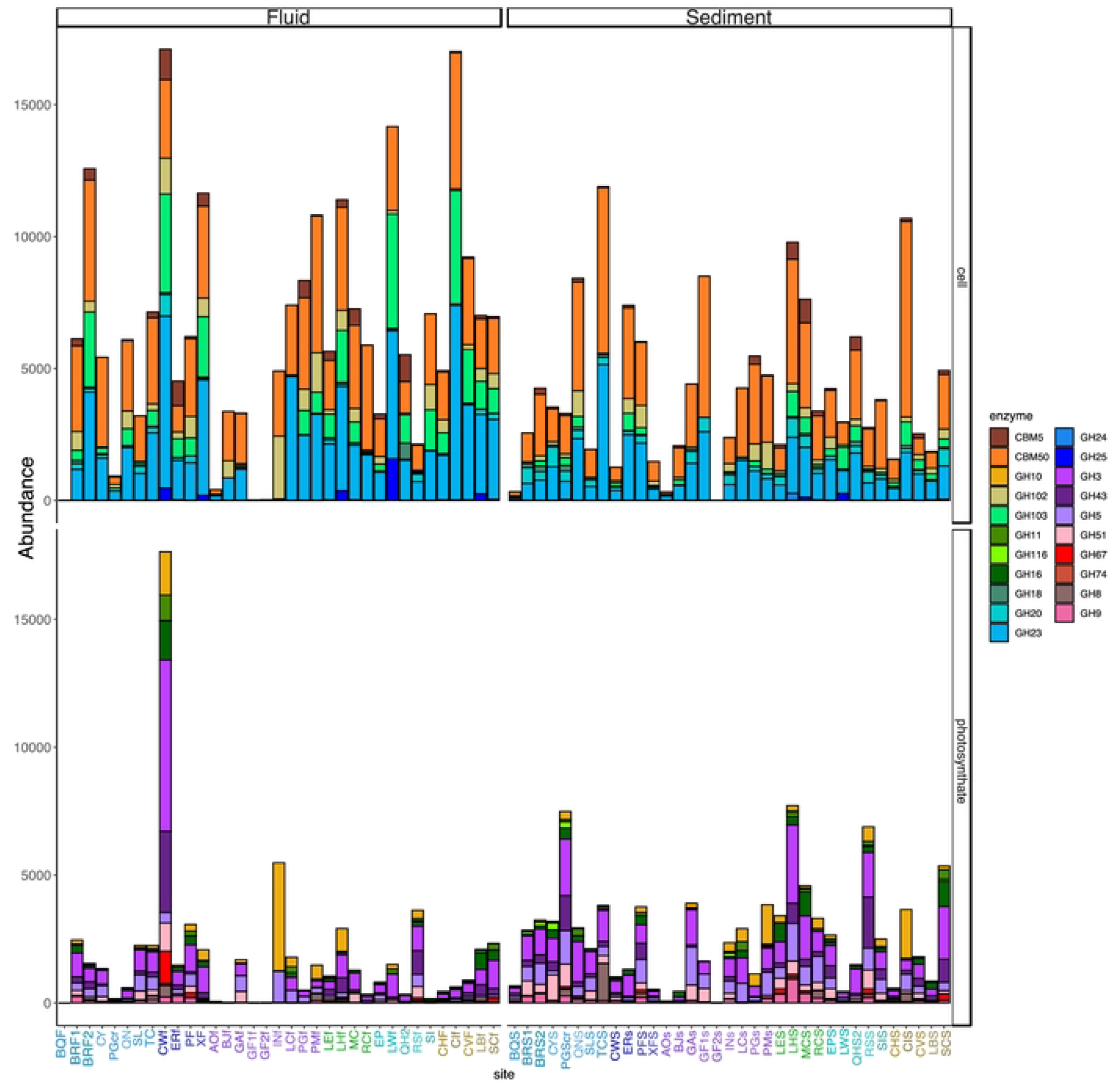
Read abundance of cell-degrading and photosynthate-degrading enzymes in metagenomic assemblies from sediments and fluids, with sites separated by geological province. Sites are colored by the geological provinces shown in Figure 1.

**Table 3:**
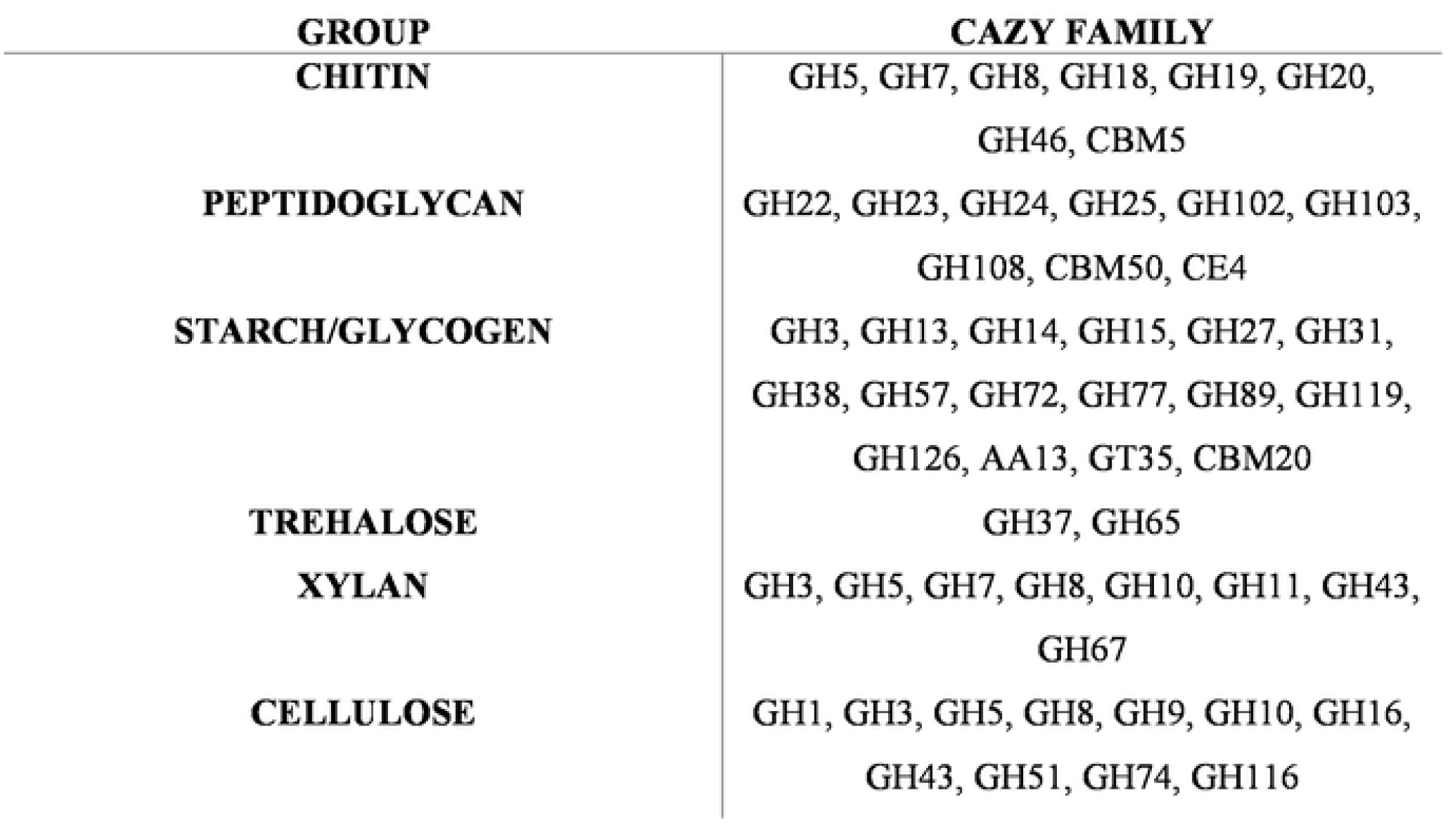
CAZy families listed based on the substrate group preference. The groups associated with cell degradation are chitin and peptidoglycan. Photosynthate degradation groups are xylan and cellulose. [48,49,60–64].

| GROUP | CAZY FAMILY |
| --- | --- |
| **CHITIN** | GH5, GH7, GH8, GH18, GH19, GH20,<br>GH46, CBM5 |
| **PEPTIDOGLYCAN** | GH22, GH23, GH24, GH25, GH102, GH103,<br>GH108, CBM50, CE4 |
| **STARCH/GLYCOGEN** | GH3, GH13, GH14, GH15, GH27, GH31,<br>GH38, GH57, GH72, GH77, GH89, GH119,<br>GH126, AA13, GT35, CBM20 |
| **TREHALOSE** | GH37, GH65 |
| **XYLAN** | GH3, GH5, GH7, GH8, GH10, GH11, GH43,<br>GH67 |
| **CELLULOSE** | GH1, GH3, GH5, GH8, GH9, GH10, GH16,<br>GH43, GH51, GH74, GH116 |

**Table 4:** ums of enzymes associated with cell and photosynthate degradation. Cell enzymes are in the chitin and peptidoglycan groups of Table 3. The photosynthate enzymes are in the groups xylan and cellulose of Table 3. The total of these two groups are summed for each site.

| SITE | CELL | PLANT | PROVINCE | TYPE | TEMP | PH | % CELL DEGRADATION |
| --- | --- | --- | --- | --- | --- | --- | --- |
| **RSS** | 2303.35 | 7311.38 | Costa Rica outer forearc | Sediment | 29.4 | 9.96 | 0.24 |
| **PGSCR** | 2717.43 | 7824.97 | Active volcanic arc | Sediment | 19.2 |  | 0.26 |
| **BQS** | 275.26 | 702.80 | Active volcanic arc | Sediment | 88.9 | 2.11 | 0.28 |
| **FAS** | 1356.01 | 3157.92 | Active volcanic arc | Sediment | 55.2 | 5.93 | 0.30 |
| **RS** | 1793.86 | 3927.14 | Costa Rica outer forearc | Fluid | 29.4 | 9.96 | 0.31 |
| **LES** | 1779.26 | 3744.37 | Cordillera Talamanca | Sediment | 34.7 |  | 0.32 |
| **BRS1** | 1790.83 | 3507.56 | Active volcanic arc | Sediment | 53.8 | 5.87 | 0.34 |
| **SLS** | 1608.47 | 2395.79 | Active volcanic arc | Sediment | 57 | 6.12 | 0.40 |
| **SCS** | 4231.40 | 5974.49 | Panama slab window | Sediment | 29.9 | 6.5 | 0.41 |
| **INS** | 1951.11 | 2651.99 | Argentina backarc | Sediment | 46.9 | 6.52 | 0.42 |
| **CYS** | 2782.04 | 3678.12 | Active volcanic arc | Sediment | 72 | 6.31 | 0.43 |
| **BR1** | 2259.54 | 2880.35 | Active volcanic arc | Fluid | 59 | 6.16 | 0.44 |
| PPS | 2768.98 | 3466.71 | Argentina backarc | Sediment | 54.3 | 8.47 | 0.44 |
| YR | 3699.46 | 4556.12 | Cordillera Talamanca | Fluid | 26 | 8.9 | 0.45 |
| RCS | 3124.49 | 3563.44 | Cordillera Talamanca | Sediment | 60 | 7.7 | 0.47 |
| GAS | 3880.17 | 4421.99 | Argentina backarc | Sediment | 67 | 6.7 | 0.47 |
| CWF | 16280.11 | 18463.25 | Active volcanic backarc | Fluid | 35 | 7.19 | 0.47 |
| INF | 4897.84 | 5482.85 | Argentina backarc | Fluid | 46.9 | 6.52 | 0.47 |
| GF2S | 8.21 | 8.62 | Argentina backarc | Sediment | 80 | 3.21 | 0.49 |
| CWS | 1102.96 | 1140.82 | Active volcanic backarc | Sediment | 35 | 7.19 | 0.49 |
| ETS | 3686.85 | 3635.11 | Active volcanic arc | Sediment | 40 | 6.06 | 0.50 |
| BRS2 | 3714.33 | 3629.83 | Active volcanic arc | Sediment | 53.8 | 5.87 | 0.51 |
| LHS | 8995.25 | 8393.51 | Cordillera Talamanca | Sediment | 55.4 | 6.7 | 0.52 |
| SL | 2882.97 | 2547.03 | Active volcanic arc | Fluid | 57 | 6.12 | 0.53 |
| PMS | 4557.13 | 3980.86 | Argentina backarc | Sediment | 50.3 | 6.53 | 0.53 |
| CVS | 2364.68 | 1971.30 | Panama slab window | Sediment | 34.9 | 7.46 | 0.55 |
| RFS | 2319.97 | 1777.15 | Argentina backarc | Sediment | 82 | 8.23 | 0.57 |
| GES | 2650.99 | 1939.12 | Cordillera Talamanca | Sediment | 35.8 | 7.8 | 0.58 |
| LCS | 4152.17 | 3007.55 | Argentina backarc | Sediment | 84 | 6.94 | 0.58 |
| PFS | 5696.03 | 4077.12 | Active volcanic backarc | Sediment | 28.7 | 5.81 | 0.58 |
| SIS | 3655.49 | 2607.84 | Costa Rica outer forearc | Sediment | 35.9 | 9.83 | 0.58 |
| MCS | 7088.09 | 5020.36 | Cordillera Talamanca | Sediment | 31.8 | 9.6 | 0.59 |
| HV21 | 5.04 | 3.52 | Spreading center hot spot | Sediment | 93.5 | 2.72 | 0.59 |
| EPS | 4095.65 | 2760.82 | Costa Rica outer forearc | Sediment | 26.4 | 9.99 | 0.60 |
| ARS | 1015.94 | 613.94 | Active volcanic arc | Sediment |  |  | 0.62 |
| PF | 5945.38 | 3326.85 | Active volcanic backarc | Fluid | 28.7 | 5.81 | 0.64 |
| GAF | 3232.45 | 1772.08 | Argentina backarc | Fluid | 67 | 6.7 | 0.65 |
| LBS | 1799.92 | 907.50 | Panama slab window | Sediment | 34.8 | 5.951 | 0.66 |
| RRF | 2427.99 | 1216.93 | Cordillera Talamanca | Fluid | 41.3 | 7.2756 | 0.67 |
| BRF1 | 5777.06 | 2784.88 | Active volcanic arc | Fluid | 59 | 6.16 | 0.67 |
| PXS | 7241.69 | 3328.99 | Active volcanic backarc | Sediment | 42.9225 | 6.3077 | 0.69 |
| XFS | 1378.69 | 617.31 | Active volcanic backarc | Sediment | 28.9 | 7 | 0.69 |
| CHS | 1497.87 | 644.82 | Panama slab window | Sediment | 31.1 | 7 | 0.70 |
| QNS | 8016.90 | 3343.38 | Active volcanic arc | Sediment | 22.9 | 5.6 | 0.71 |
| VVF | 4672.04 | 1821.79 | Argentina backarc | Fluid | 38.2 | 9.09 | 0.72 |
| SCF | 6730.07 | 2555.04 | Panama slab window | Fluid | 29.9 | 6.5 | 0.72 |
| BCF | 3840.09 | 1452.10 | Panama slab window | Fluid | 31.8 | 7.5 | 0.73 |
| CIS | 10418.56 | 3918.02 | Panama slab window | Sediment | 48.3 | 9 | 0.73 |
| BSF | 9438.61 | 3449.51 | Cordillera Talamanca | Fluid | 40.9 | 9.05 | 0.73 |
| TCS | 11507.98 | 4132.12 | Active volcanic arc | Sediment | 60 | 6.24 | 0.74 |
| ERF | 4408.88 | 1580.28 | Active volcanic backarc | Fluid | 35 | 9.51 | 0.74 |
| TC | 6889.35 | 2454.24 | Active volcanic arc | Fluid | 60 | 6.24 | 0.74 |
| BQ | 4.55 | 1.61 | Active volcanic arc | Fluid | 88.9 | 2.11 | 0.74 |
| LBF | 6797.71 | 2293.94 | Panama slab window | Fluid | 34.8 | 5.951 | 0.75 |
| QHS2 | 5764.17 | 1913.66 | Costa Rica outer forearc | Sediment | 36.7 | 8.69 | 0.75 |
| CLF | 10123.52 | 3243.36 | Panama slab window | Fluid | 50.9 | 7.5 | 0.76 |
| MTF | 7584.31 | 2352.33 | Active volcanic arc | Fluid | 59.1 | 6.32 | 0.76 |
| CY | 5257.73 | 1501.00 | Active volcanic arc | Fluid | 72 | 6.31 | 0.78 |
| TMS | 10956.68 | 3105.54 | Argentina backarc | Sediment | 69.2 | 7.13 | 0.78 |
| QHS1 | 5565.26 | 1527.88 | Costa Rica outer forearc | Sediment | 36.7 | 8.69 | 0.78 |
| GF1S | 7954.31 | 2174.13 | Argentina backarc | Sediment | 80 | 7.75 | 0.79 |
| LHF | 11274.13 | 3025.97 | Cordillera Talamanca | Fluid | 55.4 | 6.7 | 0.79 |
| EP | 3224.40 | 850.89 | Costa Rica outer forearc | Fluid | 26.4 | 9.99 | 0.79 |
| AOS | 315.56 | 82.69 | Argentina backarc | Sediment | 27.8 | 6.25 | 0.79 |
| LCF | 7355.95 | 1836.67 | Argentina backarc | Fluid | 84 | 6.94 | 0.80 |
| PGS | 5290.06 | 1307.15 | Argentina backarc | Sediment | 43.9 | 8.74 | 0.80 |
| PSS | 1165.03 | 286.77 | Costa Rica outer forearc | Sediment | 33 | 8.2 | 0.80 |
| PLS | 20079.51 | 4798.48 | Active volcanic arc | Sediment | 37.6 | 0.85 | 0.81 |
| **LEF** | 5406.90 | 1289.32 | Cordillera Talamanca | Fluid | 34.7 |  | 0.81 |
| **BJS** | 2057.82 | 474.15 | Argentina backarc | Sediment | 40 | 6.44 | 0.81 |
| **LPF** | 8018.62 | 1783.25 | Cordillera Talamanca | Fluid | 39.1 | 6.5 | 0.82 |
| **GF2F** | 25.72 | 5.37 | Argentina backarc | Fluid | 80 | 3.21 | 0.83 |
| **ERS** | 7225.99 | 1464.14 | Active volcanic backarc | Sediment | 35 | 9.51 | 0.83 |
| **QH2** | 4871.79 | 978.15 | Costa Rica outer forearc | Fluid | 48.7 | 8.53 | 0.83 |
| **HV22** | 142.26 | 28.35 | Spreading center hot spot | Sediment | 25.7 | 1.82 | 0.83 |
| **MC** | 7154.79 | 1377.19 | Cordillera Talamanca | Fluid | 31.8 | 9.6 | 0.84 |
| **XF** | 11532.69 | 2192.06 | Active volcanic backarc | Fluid | 28.9 | 7 | 0.84 |
| **PG** | 917.39 | 169.35 | Active volcanic arc | Fluid | 19.2 |  | 0.84 |
| **LWS** | 2907.58 | 491.44 | Costa Rica outer forearc | Sediment | 31.5 | 7.1 | 0.86 |
| **HAF** | 5644.82 | 936.25 | Cordillera Talamanca | Fluid | 33 | 8.9 | 0.86 |
| **BRF2** | 12377.42 | 1729.54 | Active volcanic arc | Fluid | 59 | 6.16 | 0.88 |
| **PMF** | 10782.76 | 1501.85 | Argentina backarc | Fluid | 50.3 | 6.53 | 0.88 |
| **AOF** | 399.10 | 54.82 | Argentina backarc | Fluid | 27.8 | 6.25 | 0.88 |
| **GF1F** | 12.49 | 1.69 | Argentina backarc | Fluid | 80 | 7.75 | 0.88 |
| **BW** | 3514.44 | 451.47 | Cordillera Talamanca | Fluid | 43.2 | 9.05 | 0.89 |
| **BQF** | 3.79 | 0.44 | Active volcanic arc | Fluid | 88.9 | 2.11 | 0.90 |
| **LWF** | 14067.82 | 1595.78 | Costa Rica outer forearc | Fluid | 31.5 | 7.1 | 0.90 |
| **QN** | 6016.90 | 676.83 | Active volcanic arc | Fluid | 22.9 | 5.6 | 0.90 |
| **CHF** | 4852.63 | 515.69 | Panama slab window | Fluid | 31.1 | 7 | 0.90 |
| **CV** | 9161.27 | 951.70 | Panama slab window | Fluid | 34.9 | 7.46 | 0.91 |
| **CZ** | 16531.00 | 1610.23 | Panama slab window | Fluid | 26.3 | 10 | 0.91 |
| **RC** | 5817.55 | 391.52 | Cordillera Talamanca | Fluid | 60 | 7.7 | 0.94 |
| **PGF** | 8315.79 | 515.92 | Argentina backarc | Fluid | 43.9 | 8.74 | 0.94 |
| **PBS** | 2689.73 | 112.99 | Active volcanic arc | Sediment |  |  | 0.96 |
| **CIF** | 16955.03 | 682.48 | Panama slab window | Fluid | 48.3 | 9 | 0.96 |
| **RVF** | 15451.30 | 576.05 | Active volcanic arc | Fluid | 42.7 | 6.19 | 0.96 |
| **ESF** | 7007.89 | 259.52 | Costa Rica outer forearc | Fluid | 27.9 | 9.75 | 0.96 |
| **KR22** | 61.39 | 1.58 | Spreading center hot spot | Sediment | 93 | 2.04 | 0.97 |
| **SI** | 7079.40 | 161.59 | Costa Rica outer forearc | Fluid | 35.9 | 9.83 | 0.98 |
| **BJF** | 3362.49 | 20.58 | Argentina backarc | Fluid | 40 | 6.44 | 0.99 |
| **TOTAL** | **521794.37** | **227434.98** |  |  |  |  |  |

### Multivariate analysis

We performed multivariate analysis on the datasets using a principal component analysis (PCA) to perform an unconstrained coordination analysis. For cell-degrading enzymes, sites do not cluster based on province (Figure 7A). Instead, we see that high temperature sites (HV21, HV22, KR22, GF1, GF2, and BQ) cluster together while all other sites are indistinguishable based on cell-degrading enzyme abundance. Photosynthate-degrading enzyme abundances, however, do cluster by provinces, with the Costa Rica active volcanic arc and Argentina active volcanic backarc tending to group together. Sites with volcanic activity correlate with CAZyme families GH1, GH5, GH9, GH51, and GH116, which are specifically related to cellulose degradation [48]. Here, high temperature sites continue to cluster together, but do not correlate with any specific enzymes. The nonvolcanic sites correlate with CAZy families GH16, GH43, GH74, GH11, and GH67. Families GH11, GH43, and GH67 are known to be directly related to xylan degradation [49].

**Figure 7:**
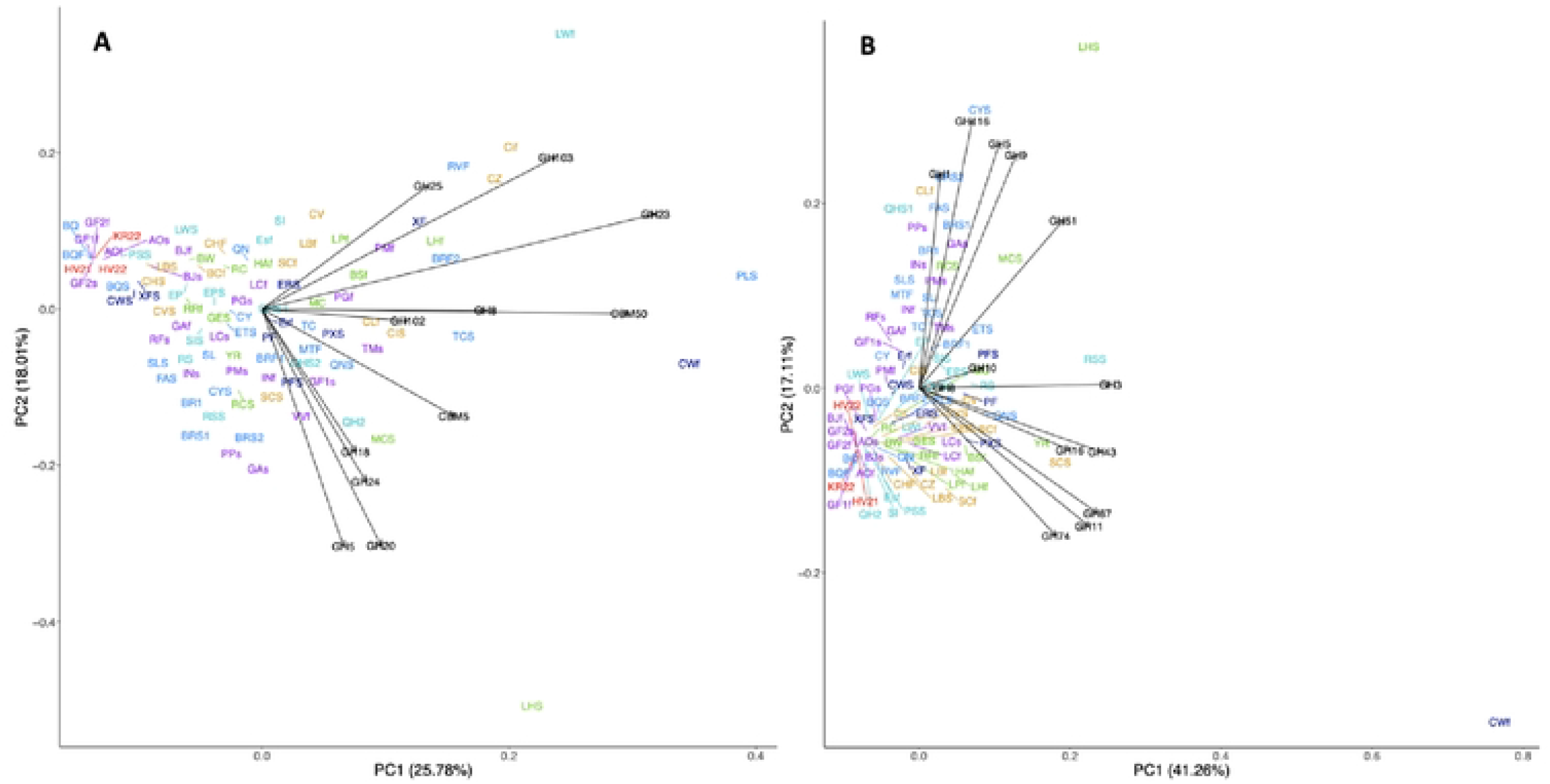
PCA plot CAZy family abundance. A. PCA plot of the abundance of CAZy families involved in the degradation of chitin and peptidoglycan, inferred to be cell-degrading. **B.** PCA plot of the abundance of CAZy families involved in the degradation of xylan and cellulose, inferred to be photosynthate-degrading. Sites are colored by the geological provinces shown in Figure 1.

To further support the idea that province separation is driven by photosynthate-degrading carbohydrate-active enzymes, rather than all peptidases, a PCA analysis was done on all the MEROPS, CAZy, and EC family annotations, as well as the annotations that correspond to the enzymes that were the target of the activity assays (Figure S2-S5). These PCA analyses show that there are no distinct clusters based on province. However, we see slight province clustering for a PCA plot of only cell-degrading and photosynthate-degrading CAZy enzymes (Figure S6).

## Discussion

All 63 sites produced a high diversity of enzymes predicted to be capable of breaking down organic matter outside the cell. This suggests that hot spring communities can break down a wide variety of organic compounds ranging from proteins and carbohydrates to structural molecules, as has been previously suggested [17,50–52]. Below we will describe how the distribution of organic carbon degrading enzymes across these sites suggest these communities are primarily supported by microbial biomass, rather than plant detritus, consistent with a chemolithoauotrophically-based ecosystem. But these heterotrophic enzymes are less directly influenced by geological features than the taxonomic compositions or respiratory capabilities of these communities.[53][54][53–55][56]

### Nature of the heterotrophic enzymes across all seeps and hot springs

We propose that the heterotrophic community in these terrestrial seeps and hot springs primarily use the organic matter of dead bacterial necromass produced *in situ* rather than allochthonous surface-derived material from plant matter. The heterotrophic community may include autotrophs that are also capable of metabolizing organic compounds. To consume complex organic matter, heterotrophs rely on extracellular enzymes such as CAZymes and peptidases to degrade the larger organic molecules so they can bring them into the cell [53]. CAZymes are important for geochemical cycling because they facilitate the breakdown of complex carbon substrates [54]. Another subset of enzymes involved in the biogeochemical cycling of heterotrophs are peptidases, which cleave peptide bonds between amino acids [47,55]. Protein degradation has been shown to be important for heterotrophs within the subsurface [56,57]. Heterotrophs within these springs may rely on protein from dead cells (necromass) for carbon and nitrogen acquisition, as they do in marine sediments [56,58]. EC annotations are based on the reactions catalyzed which allows for analysis of different reactions within the assemblies, and much of the important heterotrophic enzymes fall under EC3, hydrolases [33]. The enzyme annotation methods do not distinguish between prokaryotic and eukaryotic sources [31,33,54]. Thus, the set of enzymes present in an environment can indicate the chemical nature, and therefore the source, of organic matter being consumed by heterotrophs [59].

An ecosystem supported by bacterial and archaeal cell necromass should contain more enzymes involved in peptidoglycan and chitin degradation such as chitinases, N-acetylglucosaminidase, and lysozymes [60–63]. Enzymes that are involved in the degradation of photosynthate material, which in the case of subsurface springs would indicate allocthonous material from the surface, are often cellulases, xylanases, and glucosidases [48,49,64]. These enzymes have been characterized into CAZy families (Table 3). In our metagenomic assemblies, enzymes associated with cell necromass degradation are the most numerous overall, i.e., twice the amount of photosynthate-degrading enzyme families (Table 4). Enzymes for cell necromass consumption include those that break down peptidoglycan, a key component in bacterial cell walls. CAZyme families associated with peptidoglycan degradation are GH103, GH102, GH73, GH25, GH24, GH23, and CBM50 (Table 3) [63]. These families are found in most of the assemblies (Figure 4). The CAZyme families with the highest abundance are GH23 (peptidoglycan lyase) and CBM50 (LysM domain). These two CAZyme families are integral to the degradation of peptidoglycan.

Another facet of the heterotrophic community utilizing necromass for essential nutrients is the acquisition of starch, glycogen, and trehalose [65]. The CAZy families that are present within all assemblies are from the families GH13, GH15, GH31, GH57, GH122, and GH133 (Figure 4), which are involved in starch degradation [66]. This suggests that the possibility of using starches is a common heterotrophic metabolism across these sites.

The peptidase annotations also support the idea that the heterotrophic community relies on necromass for carbon and energy acquisition. Peptidase families such as S11, M23, S13, S66, M15, M74, M14, and C51 are used for peptidoglycan degradation (Figure 3). These families’ biological functions are associated with lysis or degradation of bacterial cell walls [60]. Of the eight listed cell wall degrading/lysing peptidases, all except M74 and C51 are present in every assembly. Peptidase family M74 is present in all but four of the assemblies, GF1f, GF1s, GF2f, and GF2s. Peptidase family C51 is present in all but eight of the assemblies, BQ, GF1f, GF1s, GF2f, GF2s, HV21, KR22, and RFs. All the samples lacking M74 and C51 fall within the temperature range (80-93.5 °C) in magmatic steam-heated springs. This may suggest that M74, which is a murein endopeptidase and C51, a D-alanyl-glycyl peptidase, are not adapted to high temperatures.

CAZyme families associated with photosynthate degradation have lower total abundance in comparison to cell degradation even though there are more possibilities for photosynthate degrading families to be found (Table 4). Of the 45 assemblies that have more than 30% photosynthate degrading enzymes out of all the enzymes related to cell and photosynthate degradation (Table 3), 34 are sediment samples (Table 4). Therefore, sediment assemblies contain more annotations for photosynthate-degrading enzymes than fluid samples. This may be caused by surface additions of photosynthetically-derived material to the sediments deposited at the surface.

The dominance of cellular biomass as the main organic matter source for the heterotrophic community is consistent with chemolithoautotrophic biomass forming the major primary production. This agrees with the findings from Barry et al., 2019a, who used helium and carbon isotopes from the same sites to show that the dissolved inorganic carbon was almost entirely derived from deep (i.e., mantle and subduction-related) sources [45]. Since hot spring enzymes related to cell necromass are more abundant in our metagenomes, we propose that hot springs microbes primarily subsist using necromass derived from chemoautotrophs rather than surficially-derived organic carbon ultimately derived from photosynthate materials.

### Independent confirmation of enzyme activity with substrate proxy analyses

Extracellular enzyme assays were used as a proxy for the activity of heterotrophic organisms within the seeps and hot spring derived sediment samples. Enzymes that are commonly-assayed in soils are often associated with cellulose and lignin degradation along with enzymes that hydrolyze proteins [59]. Degradation of plant litter is often demonstrated using assays for AG, BG, CB, and XYL, as they degrade cellulose and xylan. LEU assays represent carbon and nitrogen acquisition from proteins alongside NAG which is a proxy of the degradation of peptidoglycan and chitin [67]. Extracellular enzyme assays can reveal different organic carbon and nitrogen sources for microbial communities, allowing for inferences about biogeochemical cycling within the system. Enzymes involved in carbon and nitrogen acquisition have been used to demonstrate the limitations of nutrients within various systems [59,67,68]. The results for these assays predicted that these organisms are not highly active (Table S4). However, the fact that activity could be observed at all suggests that at least some of the enzymes we identified in our metagenomic annotations were active in natural samples and the lack of extracellular enzymatic activity may be the result of not recreating the ideal geochemical parameters for the hydrolysis within the lab setting [69].

### Highest temperature magmatic steam-heated springs have low diversity of heterotrophic enzymes

Lower diversity microbial communities tend to be found at very high temperature springs [62]. Accordingly, our high temperature and low pH sites (GF2s, GF1f, GF2f, HV22, HV21, and KR22) have fewer annotations for both CAZymes and peptidases than more mesophilic springs. This is likely due to the small number of organisms present within these sites (Figure S1). Although these sites are driven by heat and volatiles originating from the subsurface, they are not representative of deep subsurface communities. The depth of the subsurface biosphere at these sites is shallow (<50m) due to the high heat flow in the area and near boiling temperature that limit the possible distribution of microorganisms at depth [11]. The shallow subsurface nature and lower residence time of the communities in these sites combined with the lack of phylogenetic diversity is the likely cause of the lower functional diversity of the heterotrophic community.

The extracellular enzymatic assays demonstrate that many high temperature sites may not be expressing the enzymes found within their metagenomes (Table S4). The sites with the highest extracellular activities are within the temperature range of 32.5 – 50 °C. Extracellular enzymes require energy to produce, so organisms that are already spending energy surviving in very high temperature and low pH sites may not have excess capacity for extracellular enzyme production [56,70]. Therefore, because the hyperthermic sites are energy-limited they are not expressing as many extracellular enzymes as the more mesothermic springs.

### Heterotrophic metabolism is not influenced by geological processes

Chemolithoautotrophic metabolisms and the redox couples that provide power for them vary depending on geological province across the Central America convergent margin [4,7]. We hypothesized that the heterotrophic community’s enzymatic functions would also vary across the provinces, since their taxonomic identities and respiratory properties do [4,7]. However, most of the extracellular enzymes in our metagenomic assemblies do not correlate with geological provinces. Based on hierarchical clustering of spearman correlations, all sites except for the very high temperature magmatic steam-heated sites are difficult to distinguish based on abundance of predicted extracellular enzymes for peptide and carbohydrate degradation. Even though the populations of chemolithoautotrophs vary by geological setting, the biomass they produce may be similar enough that it can be broken down by similar sets of enzymes (Figure 7A), even though the heterotrophic taxa that make these enzymes also vary by geological setting [4,7].

One group of extracellular enzymes, however, does differentiate by geological province. The PCA analysis of the photosynthate-degrading enzymes shows slight clustering of the Costa Rica active volcanic arc and backarc and the Argentina backarc (Figure 7B). These sites have the same amount of photosynthate-degrading and cell-degrading enzymes as the other provinces, but the composition of the photosynthate-degrading enzymes more closely reflects cellulose degradation with enzymes such as GH5, GH9, GH116, and GH1. Springs and seeps that lack direct magmatic influence, such as the Costa Rica outer forearc, the Cordillera Talamanca, and Panama cluster separately from the volcanic sites, with more xylan-degrading enzymes. When all CAZy families for xylan, chitin, and peptidoglycan degradation are combined, clustering based on province still occurs, suggesting a strong association of volcanic sites with cellulose-degradation and non-volcanic sites with xylan-degradation (Figure S6).

Since these photosynthate-degrading enzymes are more abundant in surficial sediments than in the freshly-expressed fluids, they are likely degrading surface-derived organic matter rather than chemolithoautotrophic production in the subsurface. The surface-derived substrates that are cleaved by photosynthate-degrading enzymes, however, are unlikely to come from plants, since our sites in the Costa Rica volcanic arc are in a dense jungle and our sites from the Argentina backarc that group with them are from the high altiplano desert, which is extremely dry with little vegetation. If the photosynthate-degrading enzymes were mostly driven by introduction of surrounding plants, then the Costa Rica volcanic zone should have more similar enzymes to the other Costa Rica and Panama sites, since they are close together and have similarly dense vegetation. A more likely potential source of xylan for these springs is thermophilic algae [71,72]. Volcanic springs are often associated with higher temperatures, average 55.4°C, while the non-volcanic springs are less thermophilic with an average temperature of 37.1°C. The lower temperatures of the non-volcanic springs are closer to the optimal temperatures of growth for diverse algae, some of which are also well adapted to sulfidic and acidic sites [73]. Therefore, the non-volcanic sites may allow for algae to grow and act as a source of xylan for the heterotrophic organisms. Cellulose, which is more common in volcanic sites, is often found in cyanobacteria [74], which have a higher temperature tolerance than eukaryotic algae [75]. However, we cannot rule out the alternate possibility that these enzymes are responding to delivery of different types of subsurface-derived organic matter in the volcanic vs. non-volcanic systems.

## Conclusion

Here we present extracellular carbohydrate- and peptide-degrading enzyme potential from the metagenomes of 63 seeps and hot springs across the Central American and the Andean convergent margin (Argentinian backarc of the Central Volcanic Zone), and Iceland (mantle plume-influenced spreading center). Throughout the seven tectonic-geographic sample groups examined, we see that the heterotrophic community primarily relies on the degradation of proteins rather than carbohydrates. This is supported by the MEROPS annotations that are found in high abundance across all assemblies. The highest CAZyme and peptidase annotations are for families associated with peptidoglycan degradation. This supports the hypothesis that most of the metabolic function for heterotrophs is derived from the degradation of dead microbial cells, consistent with the major source of organic matter in this system being subsurface chemolithoautotrophic production. Very high temperature (>75°C), low pH sites (< 4), that are heated by volcanic inputs, differ from the rest of the sites based on their CAZymes and peptidases. Except for a few thermophilic peptidases, they had fewer extracellular carbon-degrading enzymes, suggesting that the secondary trophic level is less well-developed at these sites, possibly because they must put more energy into survival in these extreme conditions. Except for these high temperature sites, most extracellular CAZyme, hydrolase, and peptidase families did not differ by geological province. This suggests that, even though the taxonomic identities and respirations of the chemolithoautotrophs and heterotrophs vary by geological province [4,7], their organic matter degrading capabilities do not. The exceptions are the photosynthate-degrading enzymes which comprised a minor component of the carbon-degrading enzymes. Volcanic sites had more cyanobacteria-degrading enzymes while non-volcanic arc sites had more algae-degrading enzymes, likely due to the difference in temperature preference of those two types of phototrophs. This study revealed that the secondary community within terrestrial geothermal systems actively participates in the carbon budget within these sites by consuming chemolithoautotrophically-derived dead cell material, with enzymatic capabilities that are independent of geological province.

## Acknowledgments

The authors wish to thank Emilce Bustos, Ruben Filipovich, Joy Buongiorno, Matthew O. Schrenk, Patrick Beaudry, Bernard Marty, Diana Roman, Forrest Horton, Alan Seltzer, Maja Rasmussen, and Eemu Ranta.

## Data Availability Statement

Data is available in the NCBI SRA with project ID PRJNA627197 and project ID PRJNA914269. Iceland assemblies metagenomic workflow is available on Kbase at https://narrative.kbase.us/narrative/107494. Scripts used in R studio are available on github at rpaul5/terrestrial-geothermal-systems.

## Author Contributions

Raegan Paul^1^ Conceptualization, Data curation, Formal analysis, Investigation, Methodology, Visualization, Writing – original draft, Writing – review & editing

Timothy J. Rogers^1^ Data curation, Methodology, Resources, Software, Writing - review & editing

Kate M. Fullerton^1^ Data curation, Methodology, Resources, Software, Writing - review & editing

Matteo Selci^2^ Investigation, Writing – review & editing

Martina Cascone^2^ Data curation, Methodology, Resources, Writing – review & editing

Murray H. Stokes^1^ Investigation, Writing – review & editing

Andrew D. Steen^1^ Resources, Software, Supervision, Writing – review & editing J. Maarten de Moor3,4 Funding acquisition, Resources, Software, Supervision, Writing – review & editing

Agostina Chiodi^5^ Investigation, Writing – review & editing

Andri Stefansson^6^ Funding acquisition, Resources, Software, Supervision, Writing – review & editing

Sumundur A. Halldorsson^6^ Funding acquisition, Resources, Software, Supervision, Writing – review & editing

Carlos J. Ramirez^7^ Funding acquisition, Resources, Software, Supervision, Writing – review & editing

Gerdhard L. Jessen^8,9^ Funding acquisition, Resources, Software, Supervision, Writing – review & editing

Peter H. Barry^10^ Funding acquisition, Resources, Software, Supervision, Writing – review & editing

Angelina Cordone^2^ Investigation, Writing – review & editing

Donato Giovannelli^2,8,9,11,12,13^* Conceptualization, Funding acquisition, Methodology, Project administration, Resources, Writing – review & editing

Karen G. Lloyd^1^* Conceptualization, Funding acquisition, Methodology, Project administration, Resources, Writing – review & editing

